# Beta and theta oscillations track effort and previous reward in human basal ganglia and prefrontal cortex during decision making

**DOI:** 10.1101/2023.12.05.570285

**Authors:** Colin W. Hoy, Coralie de Hemptinne, Sarah S. Wang, Catherine J. Harmer, Mathew A. J. Apps, Masud Husain, Philip A. Starr, Simon Little

## Abstract

Choosing whether to exert effort to obtain rewards is fundamental to human motivated behavior. However, the neural dynamics underlying the evaluation of reward and effort in humans is poorly understood. Here, we investigate this with chronic intracranial recordings from prefrontal cortex (PFC) and basal ganglia (BG; subthalamic nuclei and globus pallidus) in people with Parkinson’s disease performing a decision-making task with offers that varied in levels of reward and physical effort required. This revealed dissociable neural signatures of reward and effort, with BG beta (12-20 Hz) oscillations tracking subjective effort on a single trial basis and PFC theta (4-7 Hz) signaling previous trial reward. Stimulation of PFC increased overall acceptance of offers in addition to increasing the impact of reward on choices. This work uncovers oscillatory mechanisms that guide fundamental decisions to exert effort for reward across BG and PFC, as well as supporting a causal role of PFC for such choices.

## Introduction

Accurate assessment and integration of rewards and effort into subjective value is central to healthy motivation and decision making. Reward-effort evaluation is dysfunctional across many psychiatric conditions such as depression and neurological diseases including Parkinson’s disease (PD)^1–3^. Such effects are thought to represent a transdiagnostic driver of highly disabling symptoms affecting mood and motivation, such as apathy and impulsivity^3^. Diagnosing and treating these deficits requires understanding the underlying neural signals and circuitry of motivation, which is known to be highly dependent on the interplay between prefrontal cortex (PFC) and basal ganglia (BG), under the influence of dopamine^4–6^. While a handful of studies have examined human decisions of whether to exert effort for reward using neuroimaging^7–13^, little is known about the underlying neural signals responsible. In addition, whether PFC signals are necessary for choosing to exert effort for reward, and whether PFC and BG can be differentiated in terms of their coding of effort and reward components of value, is poorly understood.

Effort-based decision-making paradigms measure motivation by offering participants different amounts of reward that require exertion of variable levels of effort^14^.

Computational modeling of these choice patterns can quantify how participants weigh and combine reward and effort into subjective value, providing the means to study the neural mechanisms of motivational states and their relationship to neuropsychiatric symptoms^15–17^. Recent frameworks for cost-benefit evaluation propose distinct roles for reward and effort computations, which may manifest differently across clinical syndromes^3,18,19^. For example, it has been proposed that apathy can be related to either insensitivity to reward or overweighting of effort costs^17,20^. Prior studies have also shown differential effects of the rates of previous rewards and efforts on choices, with the richness of recent rewards modulating exploration of options and accumulation of recent effort leading to fatigue and disengagement^13,21^.

Previous research has also identified a distributed network of brain regions underlying reward-effort trade-offs in motivated decision making. Functional MRI studies in humans have identified regions of the BG and anterior PFC, especially orbitofrontal cortex (OFC), that preferentially encode reward^13,22^, while anterior insula and dorsal subregions of medial PFC are particularly sensitive to effort^9,10,23^. However, substantial neuroimaging evidence suggests that some of these regions, including the BG and medial PFC, respond to integrated subjective value^7,8,11,24^, indicating the need for fine-grained measurements to disentangle overlapping reward and effort representations in these networks. Extensive primate studies using techniques with high spatiotemporal resolution have identified neural circuits encoding reward in the anterior PFC and OFC and BG^4,5,25–27^, but interspecies differences limit interpretability in humans^28,29^. The recent clinical availability of sensing-enabled deep brain stimulation (DBS) pacemakers for PD has created new opportunities to record local field potentials (LFPs), which reflect underlying firing rates and rhythmic synchronization within neural populations, from neural structures in awake humans^30^. Studies to date of reward coding using intracranial recordings in human PFC have mostly examined evoked potentials or high frequency activity^31–34^. To our knowledge, few have investigated low frequency, oscillatory LFP signatures of reward in human PFC^35–37^, and none have focused on motivated decision making that balances reward versus effort.

Neural oscillations have been proposed to facilitate efficient communication of choice-relevant information across distributed reward and decision networks^38,39^.

Investigations of oscillatory reward signaling in LFPs recorded from animal OFC have revealed prominent theta oscillations (4-7 Hz) that emerge at critical moments for reward learning and value-based decision making^40,41^ and are necessary for reward learning^42^. Separate lines of evidence have established that beta oscillations (12-30 Hz) in cortico-BG circuits decrease during both motor planning and execution of movement^43–46^. These beta modulations scale with movement effort^47^ and track dopaminergic states related to reward^48–50^, particularly in the low beta frequencies (12-20 Hz)^51,52^. Recent studies suggest that beta oscillations in PFC may also play an important role in effortful cognitive functions, such as attention, cognitive control, and working memory^53–57^. However, it is unclear whether PFC oscillations in theta or beta frequencies code for value, reward, or effort during effort-based decision making.

Here, we assess oscillatory signatures of reward and effort computations underlying subjective valuation in an exploratory investigation of a unique cohort of four people with PD. They were implanted with chronic PFC and BG electrodes connected to a sensing-enabled brain pacemaker as part of a separate clinical trial^58^. LFP recordings were performed during a behavioral paradigm that dissociates reward and effort components of motivated decision making^17,20^. Further, stimulation was delivered directly to PFC in one participant, providing a causal test of the role of PFC in reward-effort discounting. This arrangement facilitates dual recordings from the prefrontal cortical-basal circuit with high spatiotemporal resolution in freely moving, unconstrained people with PD outside the perioperative period, which can be confounded by surgical microlesional effects^59,60^.

We observed dissociable neural signatures of reward and effort in fronto-basal ganglia LFP recordings during decision making. PFC theta power increased according to previous trial reward, which influenced the current decision, while BG beta power decreased related to current trial effort. Furthermore, stimulation of PFC increased the number of work offers accepted and increased the positive effect of reward on decision making. These findings establish distinct oscillatory channels for processing reward and prospective physical effort demands in the PFC and BG, as well as supporting the proposal of a causal involvement of PFC in effort-based decision making.

## Results

We utilized a validated decision-making paradigm which explicitly requires accept/reject choices that trade off reward versus effort, with levels of physical effort required to obtain different levels of reward parametrically manipulated across trials (Fig. 1a). Force exertion was completed at the end of the session to avoid fatigue and movement confounds^17,20^. Four participants with chronically implanted sensing-enabled DBS devices completed the study (Fig. 1b; see Sup. Table 1 for demographics and recording details). Intracranial electrophysiological data were recorded from one pair of four contact ECoG electrodes in anterior PFC (targeted to OFC; Fig. 1c) and subcortical DBS electrodes (two participants in subthalamic nucleus (STN) and two in globus pallidus (GP)).

**Figure 1.**
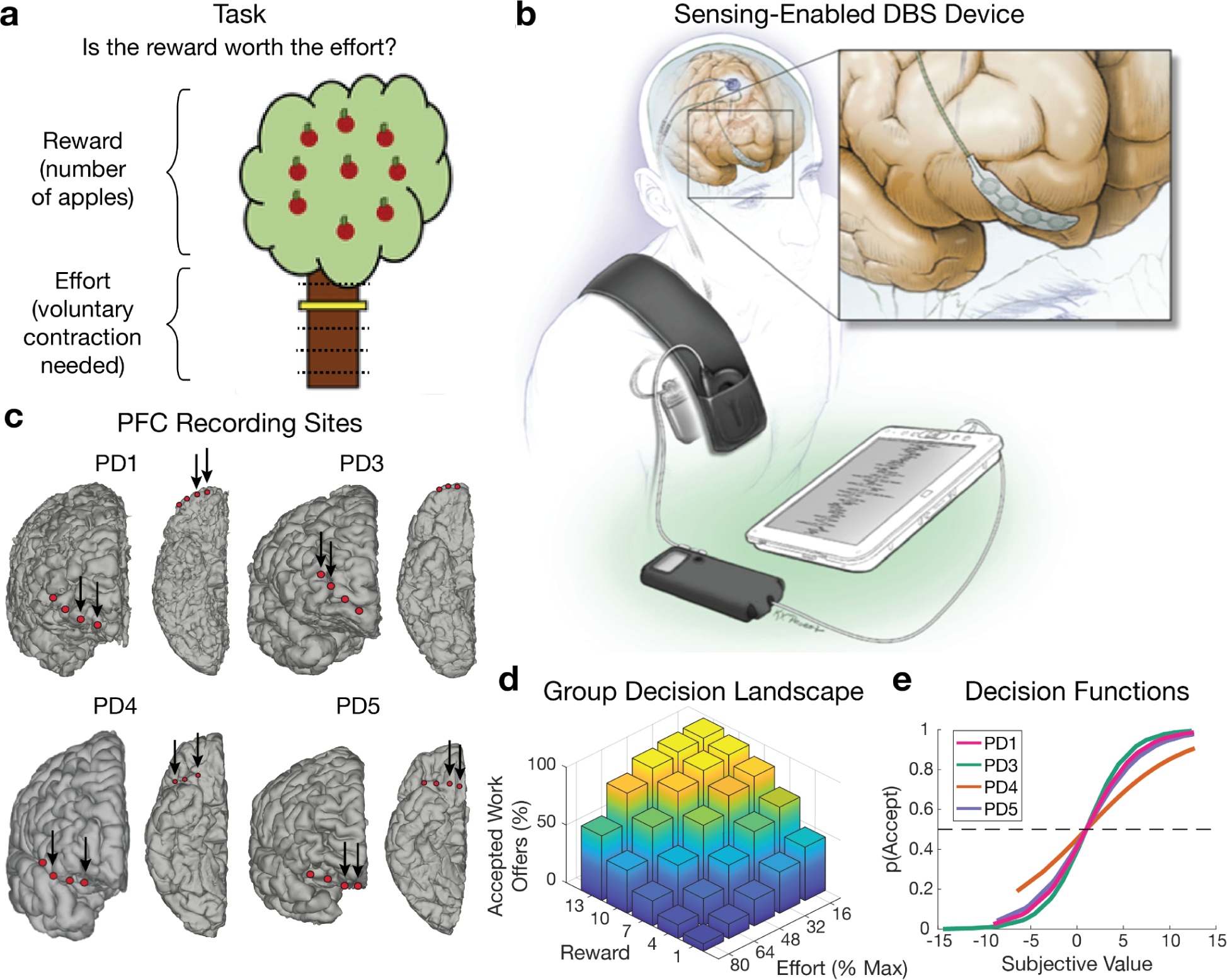
Experimental setup and behavioral results. a. Schematic of experimental paradigm showing a single trial where an offer is made of a certain number of points (apples) for a particular level of physical effort (yellow marker height) required to obtain this reward. **b** Chronically implanted recording system (Activa PC+S system shown) with wireless telemetry streaming of neural data. **c** Location of the prefrontal cortex (PFC) electrodes for each participant. Arrows indicate recording contact pairs. **d** Group-averaged performance showing percentage of accepted offers at different levels of reward and effort, confirming expected tradeoff between reward and effort. **e** Patient-specific behavioral modeling estimates the subjective value and probability of acceptance, p(Accept), for each offer.

### Behavioral performance

Participants accepted offers more often for higher rewards (*β*=0.94, *p*<10^-9^) and less often for higher effort (*β*=-2.41, *p*<10^-10^), in line with previous evidence that people devalue rewards by effort^12^ (Fig. 1d). Group-level mean reaction times (RTs) were 2.86 ± 0.73 s (Sup. Fig. 1a).

The subjective value of each offer was estimated for individual participants by fitting choice data using a previously validated model that trades off linear reward against a parabolic term capturing physical effort^12,14^. Decision policies were modeled with a softmax function to obtain p(Accept), the probability each participant would accept an offer with a given subjective value (Fig. 1e).

Generalized linear mixed models (LMMs) showed participants accepted work trials more often for offers with higher subjective value (*β*=1.808; *p*<10^-10^), and they also accepted work offers more often for easy versus hard decisions (*β*=-0.04; *p*=0.004), as measured by distance from the indifference point (i.e., p(Accept)=0.5). Participants’ RTs were faster for easier choices (i.e., high reward/low effort or low reward/high effort; *β*=-0.039, *p*<10^-4^) and faster following difficult choices on the previous trial (*β*=0.027, *p*=0.004; Sup. Fig. 1b and 1c). These results suggest participants deliberated less on easy choices and allocated more cognitive resources to decisions immediately following a difficult choice, which replicates classic within- and between-trial adaptation effects from prior studies on cognitive control^61^.

### Neural signatures of previous reward and effort

Time-frequency decompositions of the neural recordings in PFC and BG (Fig. 2a) were used to extract spectral power during the decision-making window from 0.5-1.5 seconds after offer onset, which excludes early sensory processing and subsequent motor preparation and responses (Fig. 2b and 2c; see Methods). To investigate how these regions process reward and effort during value-based decision making, we used LMMs to first test whether neural power was predicted by subjective value (integrated reward and effort), before then testing a more complex LMM with dissociable reward and effort variables. Formally, we tested which of two a priori hypotheses best predicted single-trial neural power during decision making: a simpler model composed of the subjective value of offers on the current and prior trial, and a more detailed model with separate reward and effort terms from the behavioral model for both current and previous trial offers. Previous trial predictors were included to account for effects of choice history on decision making^13,21^. These hypotheses were tested in two predefined spectral regions-of-interest in theta (4-7 Hz) and low beta (12-20 Hz), following evidence that theta is required for reward learning^42^ and that lower beta is responsive to dopaminergic manipulations^52,62^ which bias reward-effort tradeoffs^48–50^.

**Figure 2.**
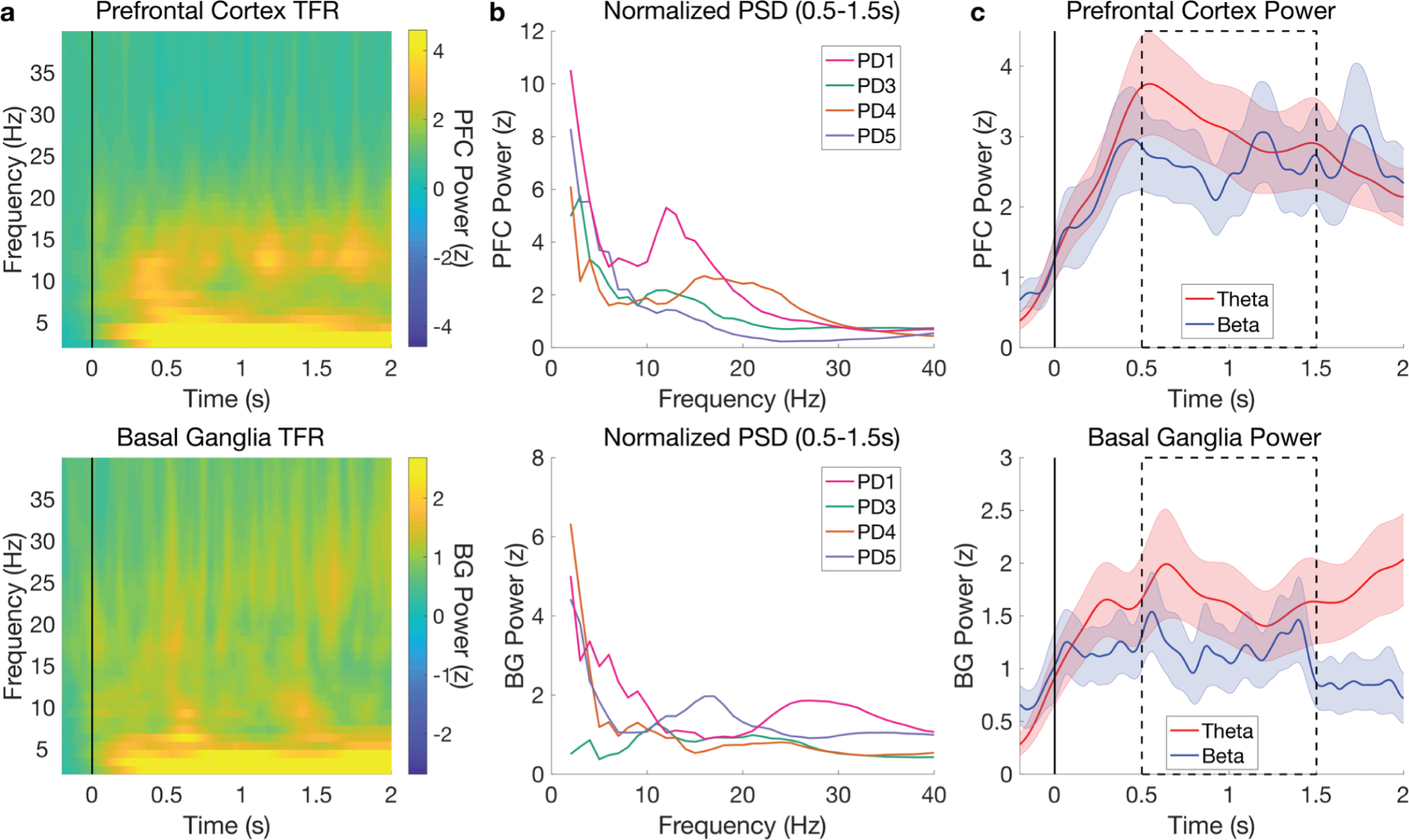
Theta and beta power in PFC and BG aligned to reward-effort stimulus presentation. a. Time-frequency representation (TFR) of group-averaged and baseline-normalized power in PFC (top) and BG (bottom), time-locked to offer presentation. **b** Baseline-normalized power in PFC (top) and BG (bottom) during decision making (averaged from 0.5-1.5 s) for each participant. **c** Group-averaged theta and beta power in PFC (top) and BG (bottom). Shading indicates standard error of the mean across participants. Dotted box indicates predefined temporal region-of-interest for single-trial modeling.

We first tested whether the subjective value model predicted spectral power of theta oscillations from 4-7 Hz (“theta power”) or spectral power of low beta oscillations (“beta power”) in both PFC and BG. Neither theta power nor beta power were significantly predicted by current or previous trial subjective value in either region (see Sup. Table 2 for full LMM results for all models). Overall, these analyses suggest that neural power in PFC and BG does not strongly represent integrated subjective value.

We then tested the second, more fine-grained model to identify dissociable reward and effort effects in theta power. We found that theta power increased with reward offered on the previous trial in PFC (*β*=0.428, p=0.017) but not BG (*β*=0.117, p=0.250), indicating that PFC theta power is sensitive to recent reward history (Fig. 3a). No other predictors reached significance (all p>0.15; Sup. Table 2).

**Figure 3.**
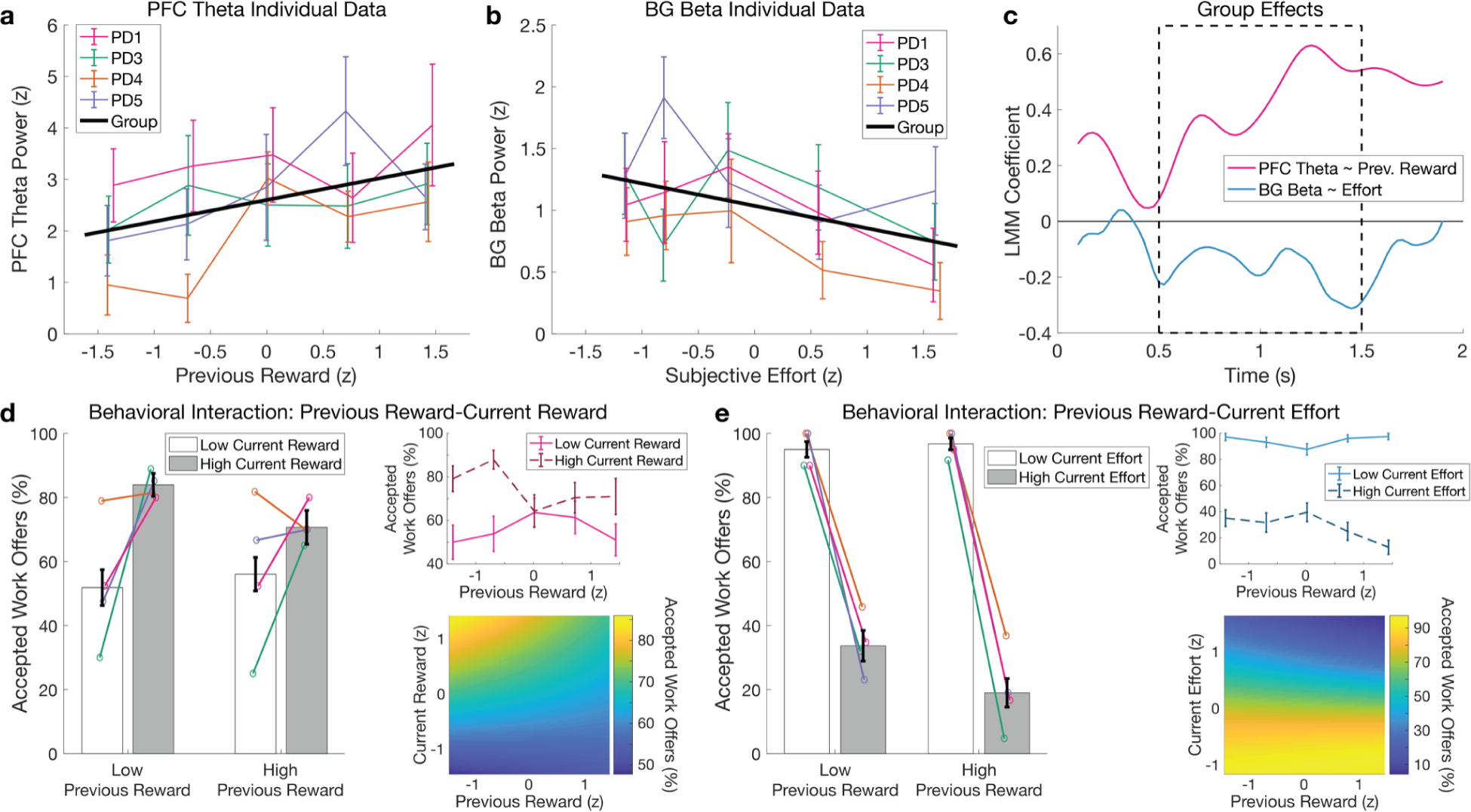
BG beta power decreases with current subjective effort, and previous reward modulates PFC theta power and choice. a. PFC theta power averaged from 0.5-1.5 s post-stimulus within each level of reward offered on the previous trial for each participant. Error bars represent standard error across trials. Black line indicates linear model fit at the group level to visualize increases in theta power with greater previous reward (*p*=0.017). **b** BG beta power decreases with greater subjective effort offered on the current trial (*p*=0.018). Plotting conventions as in **a**. **c** Time-resolved effects of previous reward on PFC theta (red) and of current subjective effort on BG beta (blue). A priori hypotheses were tested using a single model predicting data averaged from 0.5-1.5s (black dotted box). For visualization purposes only, LMMs were run at each time point to display the temporal evolution of these relationships. **d** Interaction between previous and current reward when modeling choice (*p*=0.033) plotted as the percentage of accepted offers for high and low current reward when previous reward is high or low (left). Error bars are standard error of the mean across participants, with within-subject means overlaid in the same colors as **a** and **b**. Top right inset demonstrates that the effect of current reward on the decision is greater when previous reward was low (*β*=1.189) than high (*β*=0.620) by plotting the percentage of accepted offers for high versus low reward at each level of previous reward. Bottom right inset shows partial dependence of choice on previous and current reward in the generalized LMM after controlling for other predictors. **e** The same plots as **d** visualize the interaction between previous reward and current effort when predicting choice (*p*=0.035). Current effort has a larger effect on the decision when the previous reward was high (*β*=-3.603) than low (*β*=-2.219).

Several follow up analyses were performed to clarify the role of previous reward. Behaviorally, previous reward did not reach significance in predicting the current decision using the a priori reward/effort model (*β*=-0.204, p=0.137). However, post-hoc testing of interactions between previous reward and the other predictors identified significant interactions between previous and current reward (*β*=-0.30, p=0.033) and between previous reward and current effort (*β*=-0.34, p=0.035). These interactions were examined by re-running the reward-effort model of choice separately for trials after low or high previous rewards, revealing that current rewards had a larger effect on the decision when the previous reward was low (*β*=1.189, p<10^-7^) than high (*β*=0.620, p=0.049; Fig. 3d). Conversely, the effect of current effort on the decision was greater when the previous reward was high (*β*=-3.603, p<10^-10^) than low (*β*=-2.219, p<10^-10^; Fig. 3e). Lastly, control analyses confirmed the relationship between PFC theta and previous reward could not be explained by a post-reward signal sustained from the previous trial, and PFC theta was not related to conflict as operationalized by choice difficulty (see Supplementary Information). In short, we found a positive relationship between PFC theta power and previous trial reward, and follow up analyses revealed that previous reward influenced choice by amplifying the effects of current reward and effort.

We next tested whether beta power in PFC and BG were predicted by the model with separate reward and effort predictors. We found that beta power decreased with larger subjective effort in the current offer for BG (*β*=-0.175, p=0.018; Fig. 3b) but not PFC (*β*=-0.161, p=0.211). No relationships between beta and other predictors, including reward, were significant (Sup. Table 2).

In summary, the subjective value model did not predict theta or beta power in either region, but modeling reward and effort separately revealed a dissociation of these components into distinct spectral signals across PFC and BG, with PFC theta and BG beta tracking previous reward and current effort variables, respectively (Fig. 3c). Finally, post-hoc modeling revealed that previous reward amplified the effects of current reward and effort on the decision, reflecting the behavioral relevance for the PFC theta effect.

### PFC stimulation increases accepted work offers and reward sensitivity

To test the causal role of PFC in effort-based decision making, one participant (PD5) performed the task while high-frequency stimulation was delivered to the PFC in a single-blinded, randomized, counterbalanced block-wise design (Fig. 4a). Stimulation was delivered at an amplitude below a threshold of detectability and without motor effects (see Methods). Similar to without PFC stimulation above, offers were more likely to be accepted with greater reward (*β*=2.054, *p*<10^-7^) and less likely to be accepted with greater effort (*β*=-1.274, *p*=0.002). Additionally however, there was an overall main effect of PFC stimulation, such that more offers were accepted when PFC stimulation was ON (*β*=9.763, *p*<10^-4^; Fig. 4b). There was also a significant interaction between PFC stimulation and reward (*β*=9.854, *p*<10^-4^; Fig. 4c) but not effort (*β*=-2.620, *p*=0.069; full model results in Sup. Table 3), indicating that rewards had a greater positive effect on the number of offers accepted when PFC stimulation was ON versus OFF .

**Figure 4.**
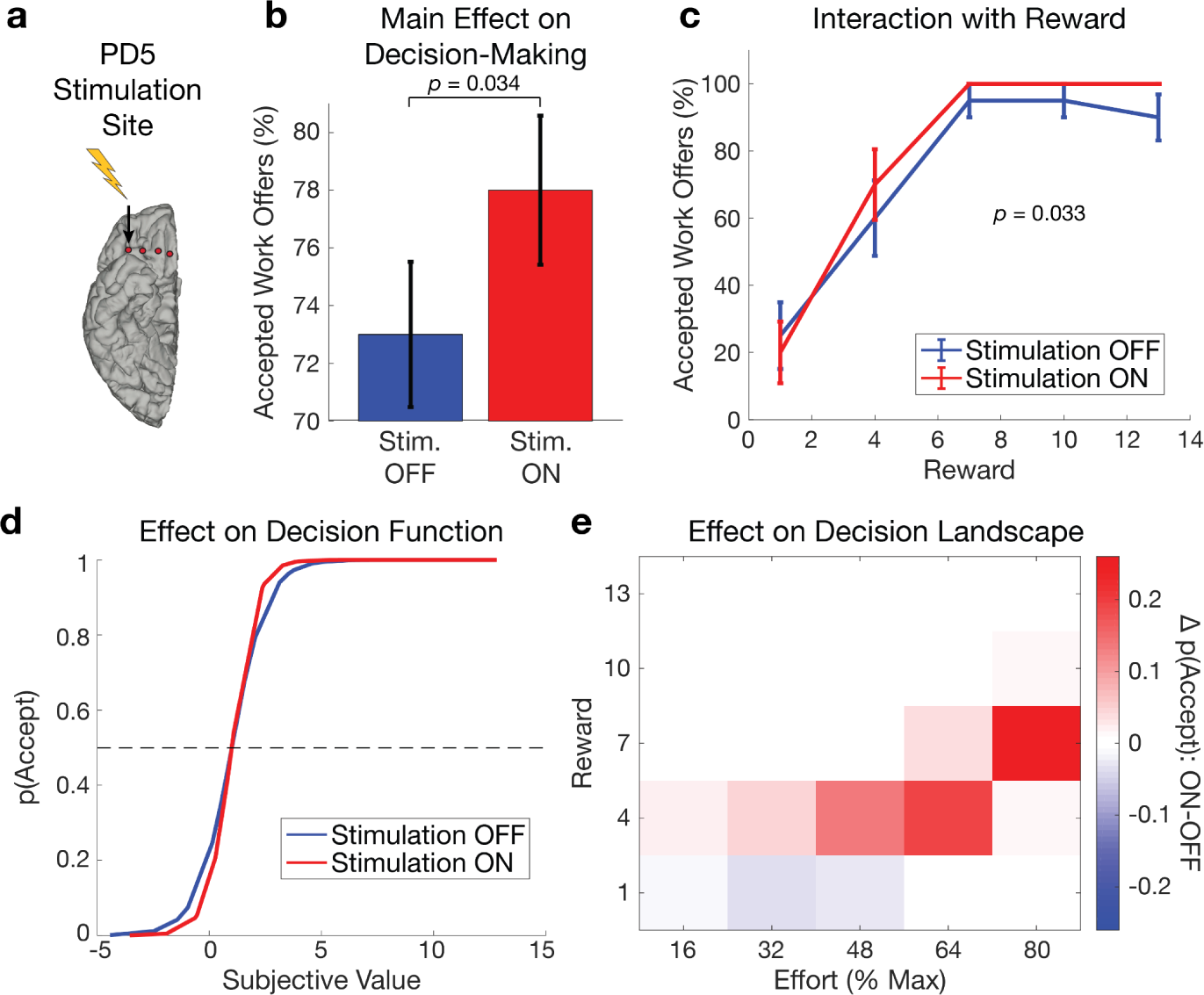
PFC stimulation increases accepted work offers and sensitivity to reward. a. Stimulation of PFC in PD5 was turned ON and OFF across blocks in a single-blind, randomized, counterbalanced order. **b** Difference in percentage of accepted work offers with PFC stimulation OFF (blue) and ON (red) indicates PD5 accepted more trials overall with PFC stimulation ON. Error bars indicate standard error across blocks. **c** Percentage of accepted work offers when PFC stimulation is ON (red) and OFF (blue) for each level of reward offered. PFC stimulation increased the positive effect of reward. Error bars indicate standard error across trials within condition, which are at ceiling for highest rewards with stimulation ON. **d** Decision functions showing the probability of accepting an offer, p(Accept), as estimated by the computational model fit separately to choice data either ON or OFF PFC stimulation. PFC stimulation decreased the randomness of choices near the indecision point (marked with dotted line). **e** Differences in p(Accept) for each combination of reward and effort when PFC stimulation is ON minus OFF. Red and blue indicate higher and lower p(Accept) when stimulation is ON, respectively. PFC stimulation primarily affected difficult choices with subjective values near the indifference point.

We also fit our behavioral model separately to choices ON and OFF stimulation. PFC stimulation increased the inverse temperature parameter of the model measuring randomness of choices, which indicates the decision policy is less random (*β*_OFF_ = 1.29; *β*_ON_ = 1.85; Fig. 4d). Plotting the difference between p(Accept) ON and OFF stimulation for each condition demonstrates that PFC stimulation appeared to have the most prominent effects on the decision function in trials with medium amounts of reward and high effort, which are challenging because they are close to the indifference point (Fig. 4e).

## Discussion

We report distinct oscillatory signatures of previous reward and effort in human PFC-BG circuits during decision making. Specifically, BG beta encoded current trial effort, and PFC theta tracked previous trial reward. Post-hoc analyses revealed that previous reward amplified the effects of current reward and effort on choice, supporting the behavioral relevance of the PFC theta signal. Spectral power of oscillatory activity in these specific frequency bands was captured by separate reward and effort predictors, and not by integrated subjective value. Finally, PFC stimulation increased the overall number of offers accepted, as well as sensitivity to reward. These results dissociate the neural basis of reward and effort computations and support a causal role of these circuits in effort-based decision making.

Our finding that theta power in PFC tracks previous reward in a reward-effort tradeoff scenario supports previous literature linking this region to reward learning and value-based decision making. Intracranial recordings from human and animal OFC have identified current and previous trial choice and reward variables encoded by local population activity^34,63^, and dopamine agonists modulate the influence of previous reward values on representations of reward expectations and prediction errors in PFC^64^. We also found that previous reward enhanced the effects of current reward and effort on decision making. This results aligns with prior evidence that it may provide context for the current decision, potentially as a direct comparison with the current reward^63^ or as a measure of recent reward history that captures the richness of the environment or global reward state^21,65^. Although the rewards on consecutive trials were independent in our laboratory paradigm, PFC likely tracks reward context because rewards have strong temporal correlations in real world environments. This neural representation of reward context information in theta may influence the current decision through interactions with high frequency or spiking activity that encodes current reward and decision variables in OFC^34,63,66^. Alternatively, recent rodent and nonhuman primate studies argue that value representations in OFC are involved in learning and do not directly affect decision making^67,68^, which suggests theta representations of reward context may instead reflect learning of the environment’s reward structure in anterior PFC^69,70^. Future studies are needed to disentangle how previous and current rewards are integrated into neural mechanisms for learning versus decision making under the influence of dopamine.

Our findings on reward information in anterior PFC theta notably contrast with a parallel literature linking theta power in dorsomedial PFC and STN to cognitive control and action monitoring in tasks eliciting sensorimotor conflict^71–75^. Our study aligns with a prior experiment recording STN LFPs in PD that also did not find low frequency conflict effects in a similar effort-based decision-making task^76^. We interpret this as demonstrating functionally distinct roles of theta oscillations in different regions (anterior versus dorsomedial PFC) and contexts (decision-making versus cognitive control paradigms), which highlights the strength of high resolution intracranial recordings for functionally segregating networks.

We have previously shown that PFC beta oscillations track depression and anxiety symptoms in the naturalistic environment in people with PD^58^. Here, we directly link beta in BG to a precise component of motivated decision making, namely prospective physical effort encoding. A prior study reported a positive relationship between frontal beta power and attentional effort and cognitive control during the delay period of a search task^53^. In contrast, we observed a negative effect of effort on BG beta during decision making in our study, which suggests the relationship between beta and effort may depend on specific cognitive and motor demands and recording sites. For example, cortico-basal ganglia beta dynamics are modulated by whether information is being encoded or cleared out of working memory^54–56^ and by whether network states are dynamic or require stabilization^77,78^. Alternatively, our finding of decreasing beta power during offer evaluation, but prior to response initiation, could be related to beta desynchronizations observed during movement planning and imagery in BG and motor networks^46,79,80^. Overall, our results support a domain-general role for beta rhythms in cognition and movement. This multiplicity of functions may help explain why dopaminergic medications and DBS, which both suppress beta^30,52,62^, affect motor and motivation symptoms in PD^81–83^.

Our dissociable reward and effort results across regions and frequency bands suggest that beta and theta during effort-based decision making are better described by independent reward and effort variables than integrated subjective value, which did not predict neural power. Previous studies have identified single units encoding subjective value in OFC, STN, and downstream regions like dorsomedial PFC that implement decisions^84–87^. This suggests the low frequency representations of reward and effort observed in our study may reflect afferent inputs to value computations performed by spiking in these regions^88^. One previous study also found that broad low frequency power (<10 Hz) in the human STN increased with reward and decreased with effort during decision making^76^. However, direct comparison to our study is hindered by differences in recording sites and task design, including their use of sequential cues for reward then effort, which has been reported to affect the valuation process^89^. For example, they also reported an integrated value signal at the second cue, which could potentially be explained by a combination of effort sensitivity with a previous reward signal akin to our finding driven by the prior reward cue. Their task also required force exertion on every trial, whereas we reserved force contractions for the end of the session. These differences indicate information presentation and temporal proximity to action may influence when and how subjective value computations unfold. Future studies recording LFPs and single units simultaneously from reward and motor networks are needed to understand how reward and effort information during decision making relate to low frequency and spiking activity across regions (e.g., PFC vs. BG, STN vs. GPi) and timescales.

Our findings dissociating neural correlates of previous reward and effort have potentially important clinical implications. Prior work in PD has shown that beta-triggered DBS in BG improves motor symptoms^81^, and DBS in BG, primarily in STN, can induce elevated mood states in people with PD^83^. Closed-loop DBS has also been trialed in depression^90^, and stimulation of OFC increased self-reported mood states in a patient with depression ^91^. In the current study, stimulation of PFC increased sensitivity to reward but not effort, matching the selectivity of PFC neural signals for reward.

Interestingly, the causal effect on decision making was largest for difficult choices when subjective value was near the indifference point. These results suggest translational potential for similar findings from non-human primates showing low current OFC stimulation increases subjective value and biases choices^92^. For example, neurodegeneration of dopaminergic circuits in PD and subsequent dopamine replacement therapy can lead to apathy or impulsivity^93,94^. If abnormal theta coding of reward context could identify deficient reward representations underlying these symptoms, such a biomarker could be used to tailor medications and neurostimulation therapies. More generally, our results suggest spectral signatures of value-based decision making in PFC and BG provide promising targets for tracking and treating motivation symptoms via closed-loop DBS in PD.

Although this dataset provides a novel opportunity to study the nature of reward-effort tradeoffs in human fronto-basal ganglia circuits, the results should be considered exploratory in nature due to the limited sample size enrolled in the parent, pilot clinical trial. Accordingly, our approach relied on strong a priori hypotheses for models and temporal and spectral analysis windows, and we are underpowered for fully unconstrained, data-driven analyses. Additionally, recordings were localized to multiple subregions of anterior PFC (orbitofrontal and frontopolar cortices) and BG (STN and GP), so future studies should also confirm the functional localization of these neural signals of reward and effort. Lastly, advances in neurotechnology that improve noise suppression techniques during stimulation will enable more sophisticated analyses of high frequency activity, phase, and network connectivity.

In conclusion, we provide evidence for the involvement of PFC-BG networks in evaluation of reward-effort tradeoffs. We identified dissociable oscillatory signatures of reward and effort in PFC theta and BG beta signals, respectively, as well as effects of single-blinded PFC stimulation on acceptance of work offers and reward sensitivity that supports the causal role of this region in reward-based decision making. The separation of reward and effort into distinct regions and frequency bands, as well as the interaction of PFC stimulation with reward but not effort, supports the segregation of these two processes. Overall, these findings constrain the role of anterior PFC and BG regions in reward learning and choice and indicate that reward and effort computations during effort-based decision making have separate neural mechanisms, which may provide more precise targets for future neuromodulatory treatments of motivation symptoms in neurological disorders.

## ACKNOWLEDGEMENTS

We would like to thank Maria Yaroshinsky for supporting data collection.

## AUTHOR CONTRIBUTIONS

Conceptualization: C.DH., M.A.J.A., M.H., P.A.S., S.L.; Methodology: C.DH., C.J.H., M.A.J.A., M.H., P.A.S., S.L.; Project Administration: S.S.W, S.J.L.; Investigation: C.DH. and S.J.L.; Formal Analysis: C.W.H. and S.J.L.; Resources: M.H., P.A.S., and S.J.L.; Supervision: M.H., P.A.S., S.J.L.; Writing - Original Draft: C.W.H. and S.J.L.; Writing - Review & Editing: C.W.H., C.DH., S.S.W., C.J.H., M.A.J.A., M.H., P.A.S., and S.J.L.; Funding Acquisition: C.W.H., C.J.H., C.DH. M.A.J.A., M.H., P.A.S., and S.J.L.

## METHODS

Resource Availability:

### Lead Contact

Further information and requests for resources should be directed to and will be fulfilled by the lead contact, Colin Hoy.

### Materials Availability

This study did not generate new unique reagents.

### Data and code availability

● Behavioral and neurophysiological data will be uploaded to a publicly available repository before publication. Raw anatomical scans will not be shared to preserve patient privacy in accordance with IRB and HIPAA regulations. Anatomical coordinates for the electrode locations will be provided in MNI group space.
● Code is publicly available in a repository on GitHub (https://github.com/hoycw/PRJ_OFC_squeeze).
● Any additional information required to reanalyze the data reported in this paper is available from the lead contact upon request.

## Experimental Model and Study Participant Details

### Human subjects approval

Patients provided written consent to participate in a protocol approved by the University of California, San Francisco (UCSF) Institutional Review Board under a physician-sponsored investigational device exemption and in accordance with the Declaration of Helsinki. The parent clinical trial under which these experiments were conducted is registered on ClincalTrials.gov (NCT03131817). Detailed descriptions of the participants and the primary study are described by de Hemptinne et al.^58^(see also Sup. Table 1). In brief, the four participants were diagnosed with idiopathic PD and mild to moderate depression and/or anxiety symptoms, but without active suicidal ideation or significant cognitive impairment, and were scheduled to undergo conventional DBS implantation for the treatment of their motor symptoms.

## Method Details

### Surgery

Participants were implanted unilaterally with quadripolor DBS leads placed in either the STN (Medtronic model 3389) or GP (Medtronic model 3387), according to clinical considerations. Placement of the DBS lead was confirmed using microelectrode recordings in the awake state^98^. In addition to the standard therapeutic DBS electrode(s) used to treat motor signs, patients were implanted with a flexible 4-contact electrocorticography (ECoG) lead (Medtronic 5387A) in the subdural space over the right PFC (Fig. 1b). ECoG contacts were 4 mm in diameter and spaced 10 mm apart.

The ECoG strips were placed over the PFC, aiming towards the orbitofrontal cortex (exact final placement shown in Fig. 1c). An intraoperative cone-beam CT merged to the preoperative MRI was used to confirm correct placement of the ECoG strip^99^. The cortical strip and ipsilateral DBS electrode were connected to lead extenders (model 37087, Medtronic), tunneled down the neck and attached to a Medtronic Activa PC+S pulse generator placed in a pocket over the pectoralis muscle under general anesthesia (Figure 1B). This investigational, bidirectional device allows both delivery of therapeutic stimulation and chronic recording of field potentials. For one patient (PD1), their Activa PC+S pulse generator was replaced at the end of battery life with the newer Medtronic RC+S device, a second generation system with improved signal-to-noise ratio for recordings during stimulation^100^.

### Electrode localization

To localize each ECoG contact in individual patients, we first used the preoperative T1 MRI to reconstruct the cortical surface using FreeSurfer^101,102^. Second, a CT scan taken 2–3 months after surgery and aligned to the T1 MRI was used to determine the location of each ECoG electrode. Each ECoG contact was projected onto the cortical surface mesh with the imgpipe toolbox^103^ using a surface vector projection method^104^.

### Experimental design

Participants performed a previously validated effort-based decision-making task^17^. The task consisted of three blocks of 25 choice trials that started with presentation of an apple tree stimulus, where the number of apples corresponded to potential point rewards (1, 4, 7, 10, or 13). A yellow mark on the trunk of the tree indicated the amount of physical effort required (16, 32, 48, 64, or 80% of maximum force) to receive the reward. Individualized maximum force was measured at the beginning of each session by asking participants to squeeze the dynamometer as strongly as possible on three trials. The maximum force exerted across those three trials was taken as the participant’s maximal voluntary contraction. On each decision trial, participants made binary choices to accept or reject the decision by pressing the left or right arrow keys on a keyboard, with their right hand. Motor mappings for the left and right choice were revealed 1.5s after the initial offer stimulus onset and randomized across trials.

Participants were given six seconds to respond. The next offer appeared after an intertrial interval with a uniform distribution between 1.6 and 1.8 seconds. To avoid fatigue influencing decisions or confounding effects of force grip preparation, participants performed actual force squeeze of a dynamometer during “point collection” trials at the end of the decision-making blocks from a random draw of previously completed trials. Participants completed three blocks of 25 decision trials per session and performed the task twice, yielding 150 total choice trials per participant. These data were collected in the laboratory while patients were on their regular dopaminergic medications and while subcortical DBS was delivered using their clinical settings.

Cortical electrodes were used for sensing. One participant (PD5; GPi) performed an additional three task sessions while receiving stimulation to the PFC. PFC stimulation was turned ON (4 blocks, 100 trials; 4 V amplitude, 130 Hz frequency, 70 µs pulse width, most lateral contact; see Fig. 4a) or OFF (4 blocks, 100 trials) across blocks within each session in a single-blinded, randomized, and counterbalanced order.

Subcortical GPi DBS was turned ON for all trials, meaning they received stimulation to both PFC and BG in ON PFC blocks, but BG stimulation only on non-PFC stimulation blocks. Neural recordings are not available from these sessions due to stimulation-related artifacts.

### Behavioral analyses

We fitted behavioral data using a computational model of subjective value developed and validated in prior studies using the same task^12,17^. Acceptance of offers was modeled on a per participant basis to determine individual subjective value (SV) using a parabolic function:

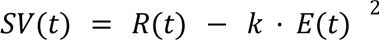

in which SV represents the subjective value on offer on trial t, R is the reward (number of apples), E is the effort level (% of maximum), and k is a free parameter that captures the participant’s subjective effort valuation for a given level of E. The participant’s decision policy was then modeled using a softmax function defined as:

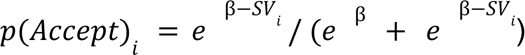

where p(Accept)*_i_* represents the probability of accepting offer i that has SV_i_, and where *β* is the temperature parameter of the softmax function which defines the stochasticity of each subject’s behavior (Fig. 1D). This decision function provides a measure of choice difficulty, which is defined as the difference in the probability of accepting an offer from the indifference point where the participant is equally likely to accept/reject the offer, i.e., p(Accept) = 0.5. Specifically, decision ease was operationalized as abs(p(Accept) - 0.5), which increases with easier and decreases with harder decisions.

### Electrophysiology recordings

PFC and BG LFPs were recorded on the PC+S device while patients performed the task. The data were then downloaded wirelessly to an external computer via bluetooth connection to the sensing-enabled DBS device. PFC recording contact selection was based on the potential therapeutic effect of cortical stimulation on mood, as studied in the clinic during a previous protocol. Briefly, stimulation was delivered from each ECoG contact sequentially while assessing mood. The contact associated with the largest improvement in subjective mood was defined as ‘potentially therapeutic’. Subsequent recordings were then performed using the contact pair with the least stimulation artifact, which was either the most distant contact pair or the contact pair surrounding the ‘therapeutic contact’. Since recordings were collected while normal clinical BG stimulation was ON, BG signals were recorded from the bipolar pair of contacts immediately surrounding the stimulation contact, which minimizes stimulation artifacts.

Neural signals in the PC+S patients were sampled at 422 Hz, with a 0.5 Hz high pass filter, and a gain of 2,000, except PD1 (RC+S) where the sampling rate was 500 Hz.

### Behavioral and electrophysiology preprocessing

Preprocessing, temporal alignment of behavioral and neural data, and analyses of behavioral and neural data used the Fieldtrip toolbox^105^ and custom code (MATLAB, Mathworks). Neural data were resampled to 1 kHz, and trials were segmented from -2 to 9 seconds relative to stimulus onset. Trials were excluded for invalid or missing responses, RTs longer than 6 seconds, and for large artifacts in the neural data, defined as containing maximum absolute values greater than two standard deviations from the mean of that trial. These behavioral and neural rejection criteria resulted in the exclusion of 2 and 16 trials for PD1, 2 and 0 trials for PD3, 2 and 2 trials for PD4, and 1 and 0 trials for PD5, respectively.

### Time-frequency representations

Spectral decompositions using the function *ft_freqanalysis* were estimated at each time point and for frequencies ranging from 2 to 40 Hz in 1 Hz steps by multiplying the signal in the frequency domain with a Hanning window of varying lengths equivalent to 4 cycles of each frequency. Task-evoked power was computed as the square of the magnitude of complex Fourier-spectra. Power values were baseline corrected by subtracting the median of the 0.8s prior to stimulus onset and dividing by the median absolute deviation of the same epoch. This median-based procedure was chosen to provide more robust control of stimulation-related noise compared to traditional mean-based z-scoring techniques^106^. Single-trial power was then computed as the average of power values within a priori determined theta (4-7 Hz) and low beta ranges (participant and electrode specific peak) from 0.5 to 1.5 seconds. This epoch was chosen to exclude initial sensory processing and preparation- and response-related motor activity once response mappings were revealed. To account for individual variability in the peak frequencies of baseline-normalized power in the low beta range (see Fig. 2b), beta power extraction in PFC and BG was customized to a 4 Hz bandwidth centered on the peak frequency for each participant and channel in the 12-20 Hz range.

## Quantification and Statistical Analysis

Linear mixed models (LMMs) were used to test whether reward, effort, and subjective value features predicted single-trial behavioral and neural data at the group level. We compared two main models to test whether behavior and neural signals were better predicted by integrated subjective value or its constituent reward and effort components. The first contained fixed effects for current and previous trial subjective value as estimated by the individual participant behavioral modeling and random intercepts for participant:

Dependent Variable ∼ SubjectiveValue_current_ + SubjectiveValue_previous_ + (1|participant)

where Dependent Variable indicates the behavioral or neural dependent variable as described below. The second model contained fixed effects for reward and subjective effort estimated by the behavioral model on the current and previous trial, as well as random intercepts for participant:

Dependent Variable ∼ Reward_current_ + Effort_current_ + Reward_previous_ + Effort_previous_ + (1|participant)

Dependent variables modeled included RTs, choices, and PFC and BG beta and theta power values. RTs were log transformed before linear modeling to account for their heavy tailed distribution, and binary accept/reject decisions were modeled using generalized LMMs with a binomial distribution. Neural dependent variables were single-trial power in theta and low beta frequencies in PFC and BG. All fixed effects predictors were z-scored within each participant. To control for outliers, trials were excluded for each model separately if the single trial values of the dependent variable exceeded three standard deviations from the mean at the group level. Of the 575 clean, preprocessed trials across all participants, this procedure resulted in exclusion of an additional 10, 12, 9, and 8 trials for modeling PFC theta, PFC beta, BG theta, and BG beta, respectively. The first trial of each block was excluded to account for weakened effects of the previous trial predictors after the rest break. Significance of fixed effects were obtained from *p* values of likelihood ratio tests testing whether the full model better described the data than a model with the fixed effect of interest removed.

Post-hoc follow up analyses to determine the role of previous reward on current choice were conducted by testing whether adding an interaction term between previous reward and current reward, current effort, or previous effort improved the reward/effort model via likelihood ratio tests. To determine the direction of the interactions, the reward-effort GLMM was run separately to predict choice for trials when the previous reward was low (1 or 4 apples) and when previous reward was high (10 or 13 apples). The data were compared between the two highest and two lowest levels of reward and effort for modeling and plotting (Fig. 3) to exclude the middle level, which would be artificially divided using a standard median split.

Since neural power is not normally distributed in our data (Lilliefors test: *p* < 10^-3^ for all power bands), we verified our LMM findings using non-parameteric permutation testing. Specifically, null distributions of *p* values were obtained from the above mentioned likelihood ratio test approach after permuting the fixed effect of interest within each subject 1000 times, and significance was measured as the proportion of null *p* values smaller than or equal to the true *p* value. All findings were the same using both approaches, so parametric LMM results are reported.

To test for canonical effects of cognitive control demands on choices and RTs^61^, we added additional fixed effects predictors for current and previous trial choice difficulty to reward-effort LMMs of behavior. Choice difficulty was defined as distance from the indifference point in the decision function, i.e., abs(p(Accept) - 0.5), which is smaller for more difficult choices. Current and previous trial choice difficulty predictors were also used in post hoc analyses of theta power to rule out alternative interpretations related to decision conflict and cognitive control (see Supplemental Information).

To visualize the temporal dynamics of our neural results, we used LMMs to predict neural power averaged in 0.2 s sliding windows stepping by 0.025 s from 0 to 2 sec after stimulus-onset. Note that statistical testing of hypotheses regarding the relationships between task variables and neural power were conducted using LMMs predicting power averaged during the pre-specified temporal and spectral regions-of-interest. The sliding time window approach is shown for visualization only.

To estimate the behavioral effect of PFC stimulation, we used a generalized linear model with a binomial distribution to predict binary choice using main effects of reward, effort, and PFC stimulation, as well as all two-way interactions. Previous trial predictors were excluded from this model because of the smaller number of trials available and greater number of main effects and interaction terms. As before, significance testing was performed using likelihood ratio tests between the full model and a reduced model without the term of interest.

## SUPPLEMENTARY INFORMATION

**Supplementary Table 1:** Patient demographics and recording details.

| Participant | PD1 | PD3 | PD4 | PD5 | Mean and SD |
| --- | --- | --- | --- | --- | --- |
| Gender | F | M | F | F |  |
| Age (years) | 53 | 55 | 67 | 70 | 61 ± 7 |
| Disease duration (years) | 25 | 5 | 6 | 9 | 11.25 ± 8 |
| Preoperative UPDRS-III (off/on) | 39/19 | 16/15 | 41/15 | 50/25 | Off: 36.5 ± 14.5<br>On: 18.5 ± 4.7 |
| Preoperative depression and anxiety severity (BDI/BAI) | 10/16 | 17/16 | 9/10 | 21/24 | BDI: 14 ± 5<br>BAI: 16.5 ± 5 |
| Preoperative cognitive state (MoCA) | 26 | 29 | 27 | 29 | 27.75 ± 1.5 |
| DBS lead side/target | R and L/STN | R/GPi | R/STN | R/GP |  |
| Coordinates of recording contact* (contact #: x,y,z) | C8: 18.6, 74.6, 6.0<br>C9: 36.9, 70.6, 1.6 | C10: 26.6, 80.3, 2.4<br>C11: 33.9, 76.6, 6.7 | C8: 13.4, 45.4, 15.7<br>C10: 31.1, 13.8, 6.9 | C8: 18.2, 42.6, 11.2<br>C9: 28.3, 42.6, 9.7 |  |

*F, Female; M, Male; UPDRS, Unified Parkinson's Disease Rating Scale; off, 12 h off PD medication; on, on regular PD medication; BDI, Beck Depression Inventory; BAI, Beck Anxiety Inventory; MoCA, Montreal Cognitive Assessment; STN, Subthalamic Nucleus; GPi, Globus Pallidus interna; R, Right; L, Left.*
*\*Contact coordinates are relative to the midcommissural point.*

**Supplementary Figure 1:**
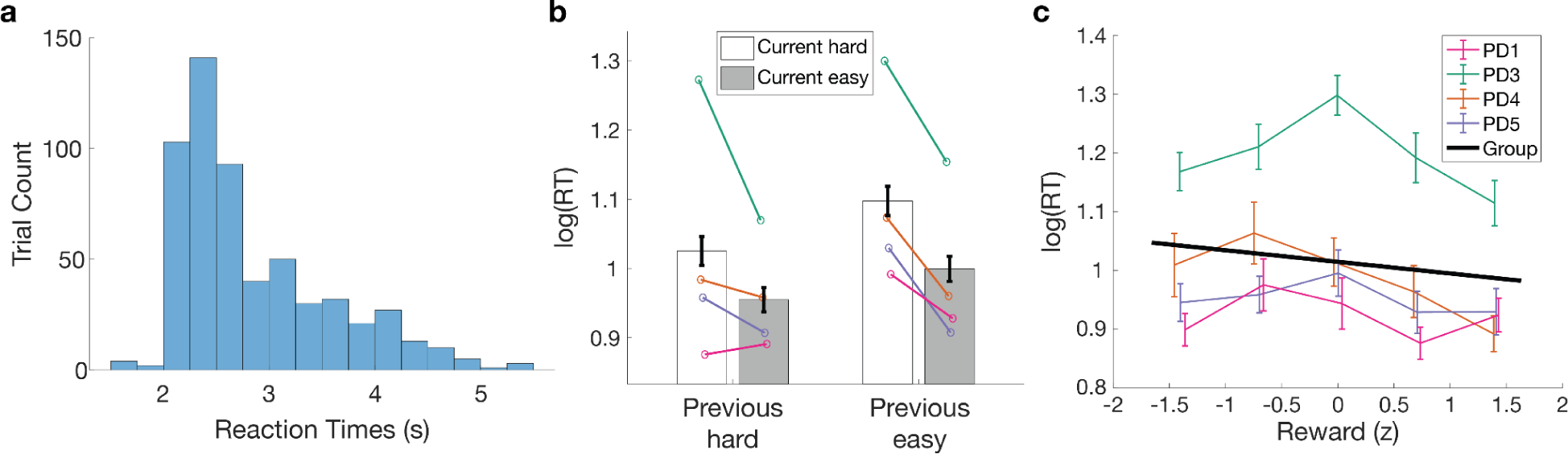
Reaction times (RTs) are sensitive to current and previous trial choice ease. **a** Distribution of RTs across participants. **b** Log-transformed RTs averaged at the group level based on median splits of current trial choice difficulty (gray for easy and white for hard) and previous trial choice difficulty (left column for hard and right column for easy). RTs were longer for difficult choices, and faster following difficult choices. Error bars indicate standard error across participants, and colored dots and lines indicate within-participant means. **c** Log-transformed RTs were significantly faster for larger rewards (*β*=-0.088, *p*=0.023) in the reward-effort model (all other *p*>0.13). Plotting conventions as in Fig. 3. However, when current and previous decision ease predictors were added to the reward-effort model, reward was no longer a significant predictor while current and previous ease effects remained significant, potentially due to correlations between reward and ease in three of four participants (*r* = 0.1, 0.76, 0.54, 0.3). Therefore, although previous studies have reported RT speeding for larger rewards (e.g., ^35^), the effect of reward on RTs in this study should be interpreted with caution because of potential collinearity concerns.

**Supplementary Table 2:**
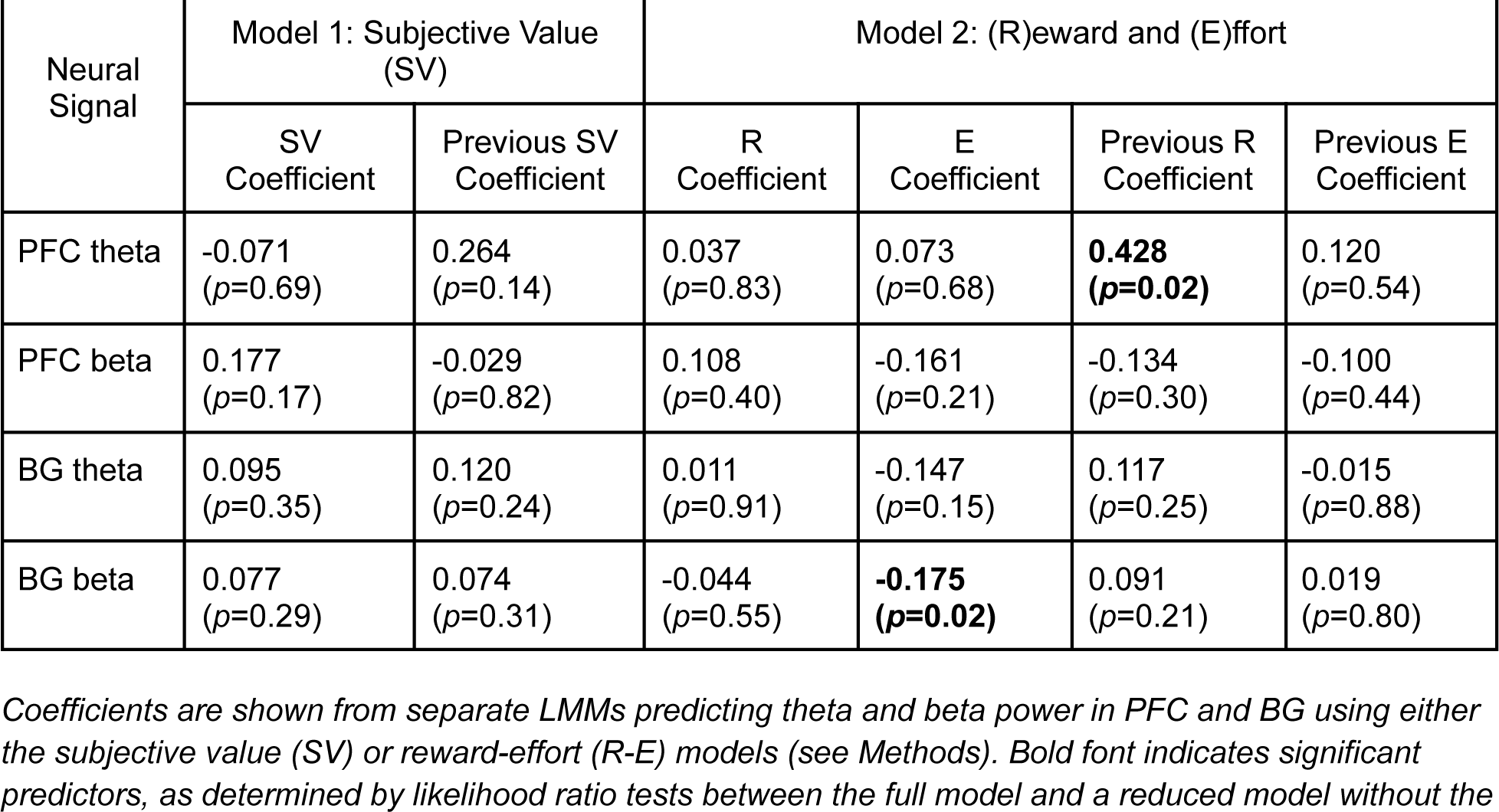
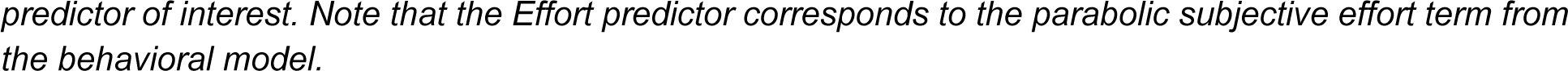
Summary of results from linear mixed modeling (LMM) of theta and beta power in PFC and BG.

| Neural Signal | Model 1: Subjective Value (SV) |  | Model 2: (R)eward and (E)ffort |  |  |  |
| --- | --- | --- | --- | --- | --- | --- |
|  | SV Coefficient | Previous SV Coefficient | R Coefficient | E Coefficient | Previous R Coefficient | Previous E Coefficient |
| PFC theta | -0.071<br>($p=0.69$) | 0.264<br>($p=0.14$) | 0.037<br>($p=0.83$) | 0.073<br>($p=0.68$) | **0.428**<br>($p=0.02$) | 0.120<br>($p=0.54$) |
| PFC beta | 0.177<br>($p=0.17$) | -0.029<br>($p=0.82$) | 0.108<br>($p=0.40$) | -0.161<br>($p=0.21$) | -0.134<br>($p=0.30$) | -0.100<br>($p=0.44$) |
| BG theta | 0.095<br>($p=0.35$) | 0.120<br>($p=0.24$) | 0.011<br>($p=0.91$) | -0.147<br>($p=0.15$) | 0.117<br>($p=0.25$) | -0.015<br>($p=0.88$) |
| BG beta | 0.077<br>($p=0.29$) | 0.074<br>($p=0.31$) | -0.044<br>($p=0.55$) | **-0.175**<br>($p=0.02$) | 0.091<br>($p=0.21$) | 0.019<br>($p=0.80$) |

Coefficients are shown from separate LMMs predicting theta and beta power in PFC and BG using either the subjective value (SV) or reward-effort (R-E) models (see Methods). Bold font indicates significant predictors, as determined by likelihood ratio tests between the full model and a reduced model without the
*predictor of interest. Note that the Effort predictor corresponds to the parabolic subjective effort term from the behavioral model.*

**Supplementary Table 3:** Summary of main effect and interaction results from general linear mixed modeling of decisions during PFC stimulation.

| Model Term | Reward | Effort | Stimulation | Reward-Effort Interaction | Reward-Stimulation Interaction | Effort-Stimulation Interaction |
| --- | --- | --- | --- | --- | --- | --- |
| Coefficient | **2.054** | **-1.274** | **9.763** | 0.744 | **9.854** | -2.620 |
| p value | **<10<sup>-7</sup>** | **0.002** | **<10<sup>-4</sup>** | 0.163 | **0.0001** | 0.069 |

*Bold font indicates significant predictors, as determined by likelihood ratio tests between the full model and a reduced model without the predictor of interest. Note that the Effort predictor corresponds to the parabolic subjective effort term from the behavioral model.*

### Supplementary analyses investigating theta power and decision conflict

Our primary result was that anterior PFC and BG theta power were predicted by previous trial reward. This finding aligns with previous work showing theta power in OFC has been linked to learning of reward values^42^ and circuit manipulations in invasive animal studies which indicate that value representations in OFC are involved in learning but do not directly influence choice^67,68^. In contrast, it has also been demonstrated that a distinct theta signal in dorsomedial PFC (dmPFC) is involved in cognitive control of decision making during actions and performance monitoring^71,72^. In particular, human intracranial studies report increased theta communication between STN and dmPFC when cognitive control is needed during difficult and high conflict decisions, as well as after errors^73,75,107,108^.

To address whether this alternative conflict framework could explain theta power in our anterior PFC and BG data, we conducted a separate analysis to test the relationships between theta power and choice difficulty, as measured by distance from this indifference point in the decision function from our behavioral model. PFC and BG theta were not predicted by choice difficulty on the current or previous trial in this task (all p>0.20), indicating our results cannot be explained by difficulty or conflict monitoring.

Lastly, we performed a post hoc control analysis to confirm the effect of previous reward on theta power was specific to the current trial. Specifically, we tested whether this signal was carried over from post-reward signals that may have been sustained from the prior trial. Modeling theta power in PFC and BG in the time period (1 second) following the decision (no feedback was delivered) on the previous trial revealed no effect of current trial reward (PFC: *β*=0.008, p=0.82; BG: *β*=0.011, p=0.77), which argues against a potential feedback-related signal spreading into the baseline normalization epoch to confound analyses on the next trial.

### Citation Diversity Statement

Recent work in several fields of science has identified a bias in citation practices such that papers from women and other minority scholars are under-cited relative to the number of such papers in the field^95–97,109^. Here we sought to proactively consider choosing references that reflect the diversity of the field in thought, form of contribution, gender, race, ethnicity, and other factors. First, we obtained the predicted gender of the first and last author of each reference by using databases that store the probability of a first name being carried by a woman^95^. By this measure and excluding self-citations to the first and last authors of our current paper), our references contain 7.62% woman(first)/woman(last), 6.67% man/woman, 26.08% woman/man, and 59.64% man/man. This method is limited in that a) names, pronouns, and social media profiles used to construct the databases may not, in every case, be indicative of gender identity and b) it cannot account for intersex, non-binary, or transgender people. Second, we obtained predicted racial/ethnic category of the first and last author of each reference by databases that store the probability of a first and last name being carried by an author of color^110,111^. By this measure (and excluding self-citations), our references contain 11.37% author of color (first)/author of color(last), 16.34% white author/author of color, 21.75% author of color/white author, and 50.54% white author/white author. This method is limited in that a) names and Florida Voter Data to make the predictions may not be indicative of racial/ethnic identity, and b) it cannot account for Indigenous and mixed-race authors, or those who may face differential biases due to the ambiguous racialization or ethnicization of their names. We look forward to future work that could help us to better understand how to support equitable practices in science.

